## Supplementary Figures for "Role of Hepatocyte RIPK1 in Maintaining Liver Homeostasis during Metabolic Challenges"

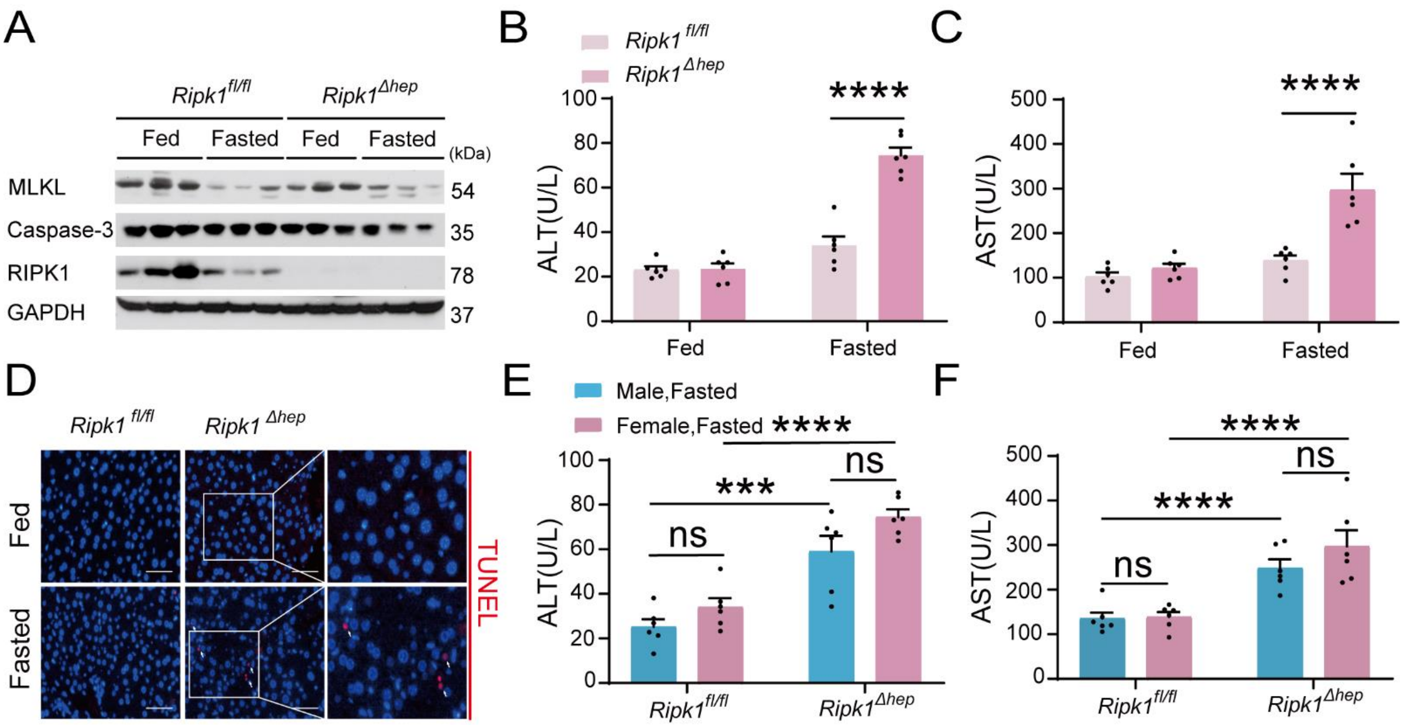


**Supplementary Figure 1.** RIPK1 deficiency in hepatocytes sensitizes the liver to short-term fasting-induced liver injury and hepatocyte apoptosis in female mice. (A) Western blot analysis of MLKL, Caspase-3, RIPK1 and GAPDH in liver tissue. (B) Serum alanine amino-transferase (ALT) levels. (C) Serum aspartate amino-transferase (AST) levels. (D) Representative fluorescence microscopy images of TUNEL staining. Scale bar, 50 μm. (E&F) There was no obvious difference between male and female mice in serum ALT (E) and AST (F) levels upon fasting. The data was analyzed via two-way ANOVA or one-way ANOVA. Data are expressed as mean ± SEM (n = 6 per group). Asterisks denote statistical significance. ns, no significant, * $P < 0.05$, *** $P < 0.001$, **** $P < 0.0001$.

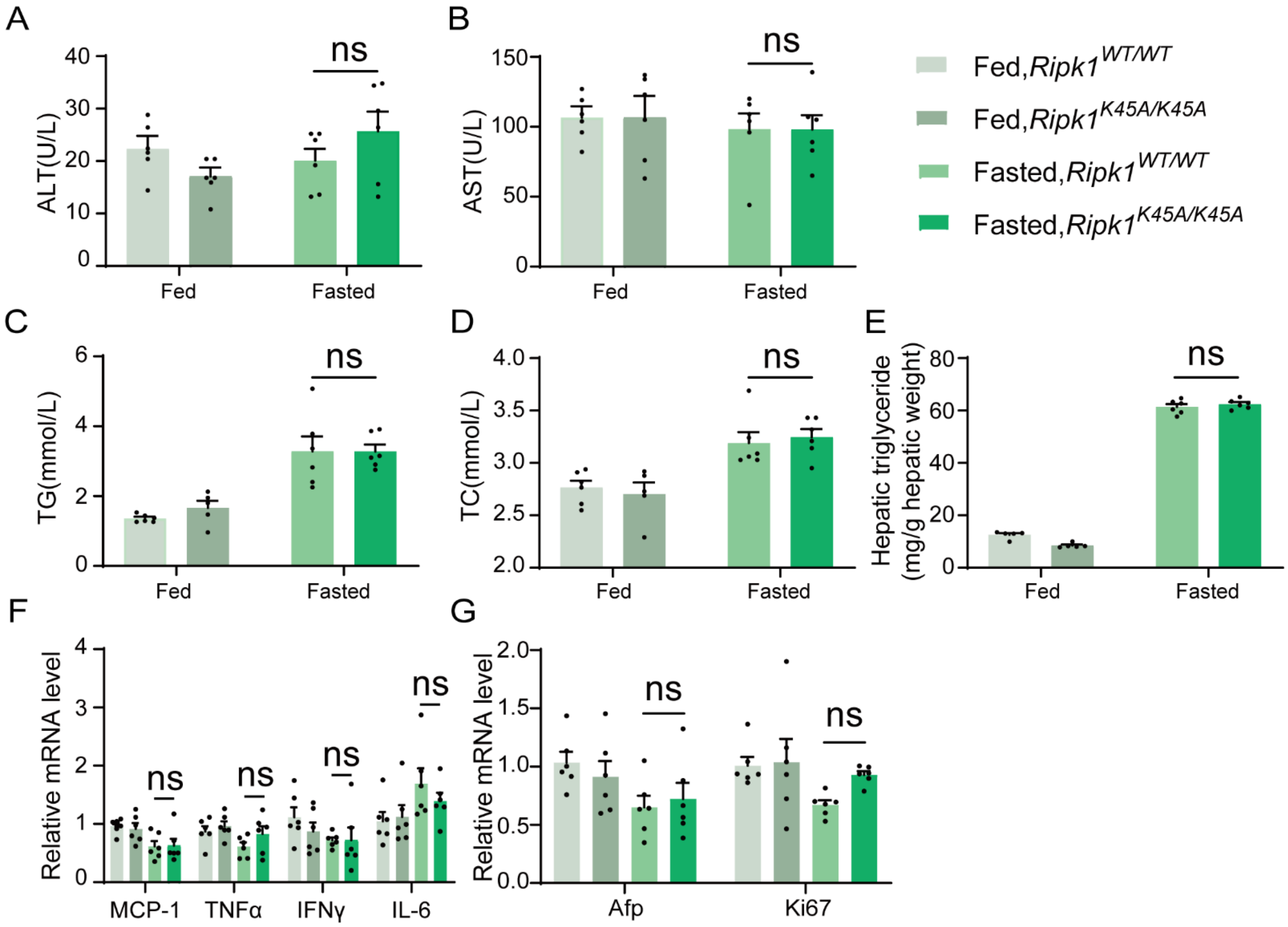


**Supplementary Figure 2.** Analysis of *Ripk1*$^{WT/WT}$ and *Ripk1*$^{K45A/K45A}$ mice before and after a 12-hour fasting period. (A) Serum alanine amino-transferase (ALT) levels. (B) Serum aspartate amino-transferase (AST) levels. (C) Serum triglycerides (TG) levels. (D) Serum total cholesterol (TC) levels. (E) Hepatic triglyceride (TG) levels (mg/g tissue). (F) Expression of inflammatory genes in the liver, assessed by qPCR. (G) Transcriptional expression of Afp and Ki67 in liver tissue. The data was analyzed via two-way ANOVA or one-way ANOVA. Data are expressed as mean ± SEM (n = 6 per group). ns, no significant, * $P < 0.05$, *** $P < 0.001$, **** $P < 0.0001$.

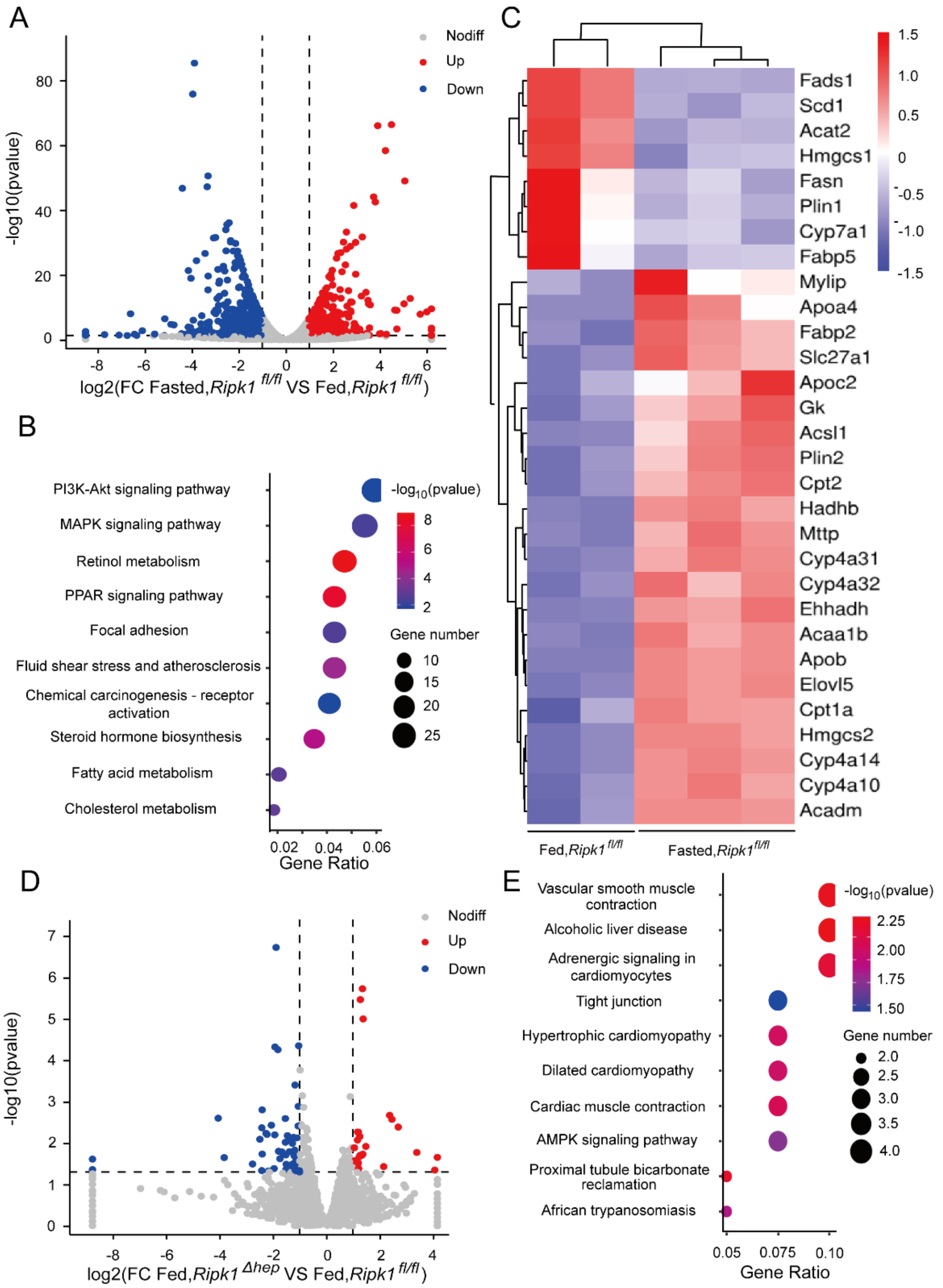


**Supplementary Figure 3.** Transcriptome sequencing of the liver tissue from *Ripk1*$^{fl/fl}$ and *Ripk1*$^{\Delta hep}$ mice. (A) The volcano plot of differentially expressed genes was illustrated. The blue spots represent the down-regulated genes in fasted group compared with control (fed) group and the red spots represent the up-regulated genes in *Ripk1*$^{fl/fl}$ mice. (B) The altered signaling pathways were enriched by KEGG analysis. (C) The genes which expression were significantly altered in fasted group were depicted in the heat map. (D) The volcano plot of differentially expressed genes was illustrated. The blue spots represent the down-regulated genes in *Ripk1*$^{\Delta hep}$ group compared with control (*Ripk1*$^{fl/fl}$) group and the red spots represent the up-regulated genes in *Ripk1*$^{\Delta hep}$ group. (E) The altered signaling pathways were enriched by KEGG analysis.

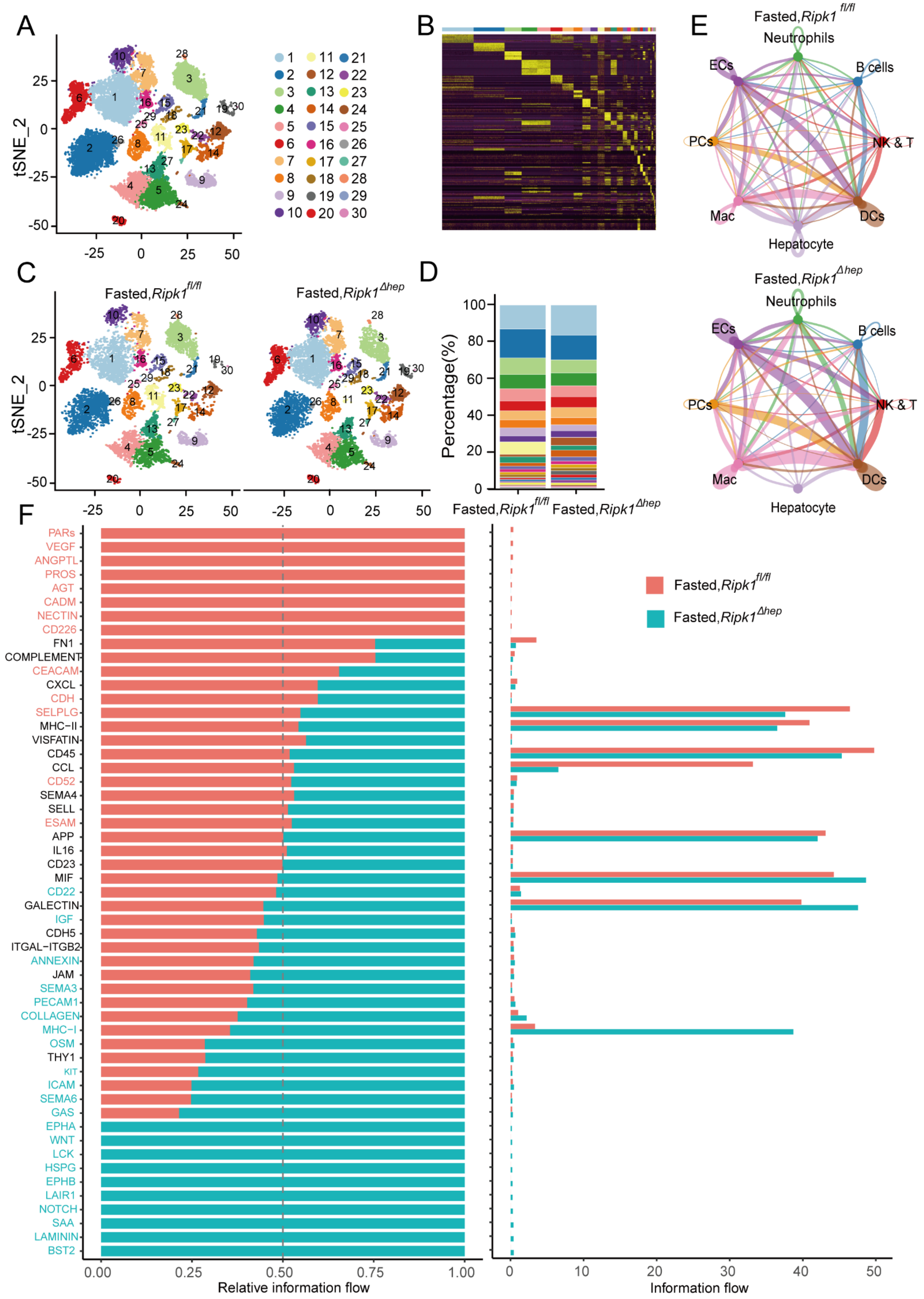
A
tSNE_2
B
C
Fasted,Ripk1 fl/fl
Fasted,Ripk1 Δhep
D
Percentage(%)
E
Neutrophils
B cells
NK & T
DCs
Hepatocyte
Mac
PCs
ECs
F
PARs
VEGF
ANGPTL
PROS
AGT
CADM
NECTIN
CD226
FN1
COMPLEMENT
CEACAM
CXCL
CDH
SELPLG
MHC−II
VISFATIN
CD45
CCL
CD52
SEMA4
SELL
ESAM
APP
IL16
CD23
MIF
CD22
GALECTIN
IGF
CDH5
ITGAL−ITGB2
ANNEXIN
JAM
SEMA3
PECAM1
COLLAGEN
MHC−I
OSM
THY1
KIT
ICAM
SEMA6
GAS
EPHA
WNT
LCK
HSPG
EPHB
LAIR1
NOTCH
SAA
LAMININ
BST2
Relative information flow
Information flow

**Supplementary Figure 4.** Single-cell RNA sequencing of liver tissue from *Ripk1*$^{fl/fl}$ and *Ripk1*$^{\Delta hep}$ mice. (A) t-SNE visualization depicting the clustering of liver cells based on 22,274 single-cell transcriptomes. (B) Heatmap illustrating the marker genes associated with each cluster. (C) t-SNE plots displaying color-coded cell clusters of cells in the *Ripk1*$^{fl/fl}$ (left) and *Ripk1*$^{\Delta hep}$ (right) mice liver tissues. (D) Bar charts showing the proportion of major cell clusters among all cells at different genotypes after fasting by scRNA-seq. (E) Circle plots displaying putative ligand-receptor interactions among eight cell subtypes in the liver tissue of *Ripk1*$^{fl/fl}$ (top) and *Ripk1*$^{\Delta hep}$ (bottom) mice. (F) All significant signaling pathways were ranked based on their differences in overall information flow within the inferred networks between *Ripk1*$^{fl/fl}$ and *Ripk1*$^{\Delta hep}$ mice. The overall information flow of a signaling network was calculated by summing all the communication probabilities within that network.
